# Deep learning based behavioral analysis enables high precision rodent tracking and is capable of outperforming commercial solutions

**DOI:** 10.1101/2020.01.21.913624

**Authors:** Oliver Sturman, Lukas von Ziegler, Christa Schläppi, Furkan Akyol, Benjamin Grewe, Johannes Bohacek

## Abstract

To study brain function, preclinical research relies heavily on animal monitoring and the subsequent analyses of behavior. Commercial platforms have enabled semi high-throughput behavioral analyses by providing accurate tracking of animals, yet they often struggle with the analysis of ethologically relevant behaviors and lack the flexibility to adapt to variable testing environments. In the last couple of years, substantial advances in deep learning and machine vision have given researchers the ability to take behavioral analysis entirely into their own hands. Here, we directly compare the performance of commercially available platforms (Ethovision XT14, Noldus; TSE Multi Conditioning System, TSE Systems) to cross-verified human annotation. To this end, we provide a set of videos - carefully annotated by several human raters - of three widely used behavioral tests (open field, elevated plus maze, forced swim test). Using these data, we show that by combining deep learning-based motion tracking (DeepLabCut) with simple post-analysis, we can track animals in a range of classic behavioral tests at similar or even greater accuracy than commercial behavioral solutions. In addition, we integrate the tracking data from DeepLabCut with post analysis supervised machine learning approaches. This combination allows us to score ethologically relevant behaviors with similar accuracy to humans, the current gold standard, thus outperforming commercial solutions. Moreover, the resulting machine learning approach eliminates variation both within and between human annotators. In summary, our approach helps to improve the quality and accuracy of behavioral data, outperforming commercial systems at a fraction of the cost.

## Introduction

Accurate analysis of rodent behavior is of the utmost importance when assessing treatment efficacy in preclinical research. The rapid development of new tools and molecular interventions in rodents, as well as the ever increasing number of available transgenic mouse lines, increase the need to accurately and efficiently detect and quantify rodent behavior [1–3]. Typically, behavioral analysis relies on commercial equipment to track an animal’s path of movement or measure the time spent in specific areas of testing arenas. Commercial solutions usually use video tracking or infra-red beam grids, and are available either as stand-alone software packages (Ethovision, Anymaze), or are integrated with hardware to create all-in-one behavioral analysis apparatus (e.g. TSE Systems, Campden Instruments, Med Associates). Such systems have enabled researchers to conduct semi high-throughput behavioral screening [4,5]. However, these commercial solutions are not only very expensive, but also lack the ability to flexibly define and score specific behaviors of interest. Further, they cannot be easily adapted to fit changing experimental needs. Even more problematically, they often measure ethological behaviors with very poor sensitivity. As a result, human scoring has remained the gold standard when scoring ethological behaviors. However, human annotation is not only excessively time consuming, but also hampered by high intra- and inter-rater variability. For instance, human annotators tire when performing repetitive tasks and their performance may vary, not only between days but also hour to hour [6]. Additionally, the complexity of animal behavior can overwhelm the annotator, and subtle differences in the definition of complex behaviors can further increase the variability between human annotators, leading to high inter-rater reliability [7,8].

Recently, major advances in machine learning have given rise to the first descriptions of unsupervised analyses of behavior, which reveal the stunning temporal and structural complexity of rodent behavior [9–13]. However, these advanced analyses are challenging for many biology and behavioral research labs to establish, which probably explains why they have not yet been widely implemented by the behavioral research community. An elegant and accessible implementation of deep learning for motion tracking and pose estimation is DeepLabCut (DLC), an open source software package that has been rapidly disseminating across laboratories throughout the world [14,15]. In contrast to commercial systems, DLC allows the user to define and track specific points of interest (e.g. specific body parts). Due to this increased level of detail and flexibility, we tested if DLC could be harnessed to replace existing commercial tracking packages, and whether it could be used to help reach human accuracy when scoring complex, ethological behaviors. Behavior tracking and analysis is performed in a vast number of behavioral tests for rodents. In this report, we focus on three of the most popular behavioral assays routinely used in preclinical research: the open field test [16,17]; the elevated plus maze [18,19]; and the forced swim test [20–22]. A search on pubmed showed that these tests have been used in more than 10’000 research papers to date, with a steady increase over the last decade **(Figure S1)**. Despite the fact that the amount of information that can potentially be gathered from the observation of behavior in these tests is almost limitless [9,10], several task-specific ethological behaviors have been identified [23,24]. Head dipping in the elevated plus maze [19,25]; rearing in the open field test [26–29]; and floating in the forced swim test [30,31] are just three examples of ethological behaviors associated with emotional and disease states [32,33]. For example, reduced exploration (rearing/head dipping) indicates anxiety [29], and floating in the forced swim test has been linked to adaptive stress-coping behaviors [34,35], although it is also frequently used to screen the antidepressant activity of new drugs [36,37]. Therefore, being able to accurately score and report these behaviors adds an important layer of information to the basic motion path of the animal. In this work we carefully compare our DLC-based approach to commercial platforms (the video tracking software EthoVision XT14 from Noldus, and the ‘all-in-one’ Multi Conditioning System from TSE systems), and to behavior rated by several human annotators (the gold standard). This comparison is valuable for the field as a thorough assessment of the reliability of different commercial platforms is currently lacking.

## Methods

### Animals

C57BL/6J (C57BL/6JRj) mice (male, 2.5 months of age) were obtained from Janvier (France). Mice were maintained in a temperature- and humidity-controlled facility on a 12 hour reversed light–dark cycle (lights on at 08:15 am) in individually ventilated cages (SealSafe PLUS, Tecniplast, Germany) with food (M/R Haltung Extrudat, Provimi Kliba SA, Switzerland, Cat.# 3436) and water ad libitum. Cages contained wood chip bedding (LIGNOCEL SELECT, J. Rettenmaier & Söhne, Germany) nesting material (tissue paper) and a transparent red plastic shelter. Mice were housed in groups of 5 per-cage and used for experiments when 2.5-4 months old. All mice were given a minimum of 2 weeks to acclimatize to the light cycle and environmental conditions before testing. For each experiment, mice of the same age were used in all experimental groups to rule out confounding effects of age. All tests were conducted during the animals’ active (dark) phase from 12-5 pm. Mice were single housed 24 hours before behavioral testing in order to standardize their environment and avoid disturbing cagemates during testing [38,39]. All procedures were carried out in accordance to Swiss cantonal regulations for animal experimentation and were approved under license 155/2015.

### Open Field Test (OFT)

Open-field testing took place inside sound insulated, ventilated multi-conditioning chambers (TSE Systems Ltd, Germany). The open field arena (45 cm × 45 cm × 40 cm [L × W × H]) consisted of four transparent Plexiglas walls and a light grey PVC floor. Animals were tested under four equally spaced yellow lights (4 lux across the floor of the open field) with 65 dB of white noise playing through the speakers of each box. An infrared light illuminated the boxes so that an infrared camera could be used to record the tests. Prior to testing each animal, the entire open field arena was cleaned using 10 ml/l detergent (For, Dr. Schnell AG). The room housing the multi-conditioning chambers was illuminated with red LED lights (637 nm). Animals were removed from their homecage by the tail and placed directly into the center of the open field. The doors of the conditioning chamber were then swiftly closed. Tracking/recording was initiated by the Multi Conditioning System upon first locomotion grid beam break, whereas videos of this test were analysed from the time the doors of the box were closed (approx. 3-5 s after first beam break). All open field tests were 10 minutes in duration. Distance, time in center, supported rears and unsupported rears were recorded in the OFT.

### Elevated Plus Maze (EPM)

The elevated plus maze was made from grey PVC, with arms measuring 65.5 cm × 5.5 cm (L × W), elevated 61.5 cm. Prior to testing each animal, the entire elevated plus maze was cleaned using 70% EtOH in H_2_O. The room housing the elevated plus maze was lit with two small lamps attached to the ceiling, they were adjusted until the open arms were at approximately 19-21 lux. A blackout curtain separated the room so that light from the screen would not alter the light conditions in the room and so the rater could not be seen by the animal. Animals were removed from their homecage by the tail and placed directly into the center of the EPM using a small starting box. Tracking/recording was initiated automatically by Ethovision XT14 (upon start condition: center point in arena for 2 seconds) and at the beginning of the video in DeepLabCut. All elevated plus maze tests were 10 minutes in duration. Distance, velocity, time in zone (open/closed arms + center) and head dips were recorded in the EPM.

### Forced Swim Test (FST)

Animals were moved from the colony room to a holding room before immediate forced swim testing in 17.9-18.1°C water for 6 minutes. The forced swim took place in a plastic beaker (20 cm diameter, 25 cm deep, filled to 17 cm so no mouse could touch the bottom of the container with its tail, or escape). Tracking/recording was automatically initiated by Ethovision XT14 as the mouse made contact with the water. The beaker was cleaned and the water was changed and shortly before each swim. Overhead red LED lights (637 nm, invisible to the mice) dimly illuminated both the holding and testing rooms. Infrared LED strips illuminated a white POM-C box onto which the beaker was placed. Distance, velocity and floating were recorded in the FST.

### Noldus EthoVision

Ethovision XT14 was used to acquire all forced swim and elevated plus maze videos and to analyse all of the open field videos. The automatic animal detection settings were used for all tests, slight tuning of these settings was performed using the fine-tuning slider in the automated animal detection settings to ensure the animals could be tracked throughout the entire arena. We ensured there was a smooth tracking curve and that the center point of the animal remained stable before analysis took place.

### DeepLabCut (DLC)

DeepLabCut 2.0.7 was used to track all points of interest **(Figure 1)**. The OFT network was trained using 15 frames from 8 randomly selected videos for 1030000 iterations (multistep: 0.005 (10000 iterations), multistep: 0.02 (430000 iterations), multistep: 0.002 (730000 iterations), multistep: 0.001(1030000 iterations). The EPM network was trained using 10 frames from 13 randomly selected videos for 250000 iterations (multistep: 0.005 (2500 iterations), multistep: 0.02 (12500 iterations), multistep: 0.002 (187500 iterations), multistep: 0.001(250000 iterations). 10 outlier frames from each of the training videos were then corrected, with points with a p < 0.7 being relabelled. The network was then refined using the same number of iterations. The FST network was trained using 20 frames from 28 randomly selected videos for 250000 iterations (multistep: 0.005 (20000 iterations), multistep: 0.02 (100000 iterations), multistep: 0.002 (175000 iterations), multistep: 0.001(250000 iterations). 20 outlier frames from each of the training videos were then corrected, with points with a p < 0.7 being relabelled. The network was then refined using the same number of iterations). Although we provide the exact number of training iterations/frames, we observed that exact number of training iterations/frames and refined outlier frames is not particularly important, since the networks worked similarly with slightly different numbers of iterations/frames. The data generated by DeepLabCut was then processed using custom R Scripts that are available in the supplementary data (https://github.com/ETHZ-INS/DLCAnalyzer).

**Figure 1.**
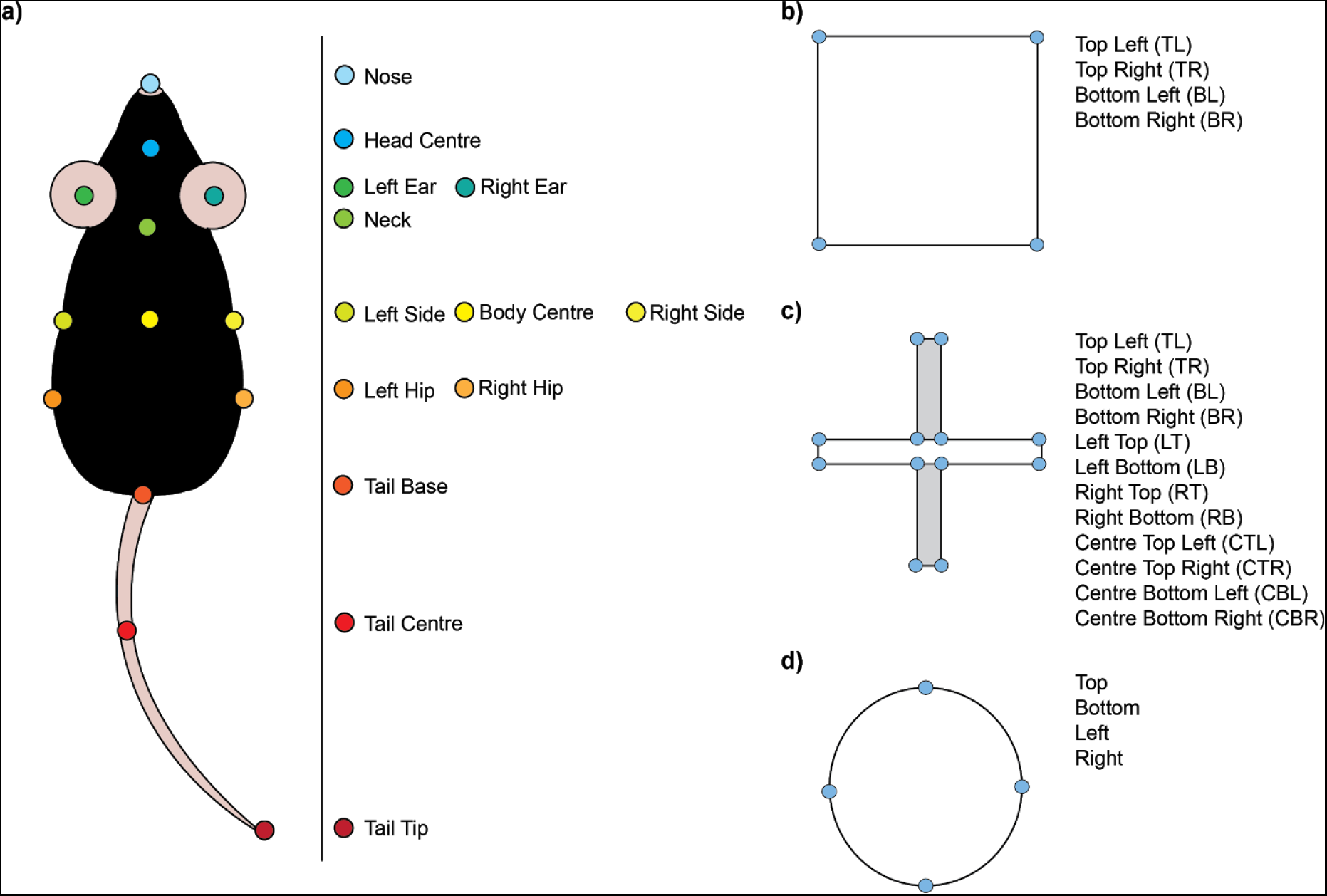
The labels used to train our DLC networks. **(a)** The standardised points of interest used to track the animal. **(b,c,d)** The points of interest required to track the animal in the open field **(b)**, the elevated plus maze **(c)** and the forced swim test **(d)**.

### TSE Multi Conditioning System

Locomotion was tracked using an infrared beam grid; an additional beam grid was raised 6.5 cm above the locomotion grid to measure rearing. The central 50% (1012.5 cm^2^) was defined as the center of the arena. To automatically distinguish supported from unsupported rears, we empirically determined the area in which mice could not perform a supported rear. Thus, all rears within 12.5 cm of the walls were considered supported rears, while rears in the rest of the field were considered unsupported rears. Rearing was defined as an interruption of a beam in the z-axis for a minimum of 150 ms. If another rear was reported within 150 ms of the initial rear, it was counted as part of the initial rear.

### Analysis of DLC Coordinates

X and Y coordinates of tracked points as determined with DLC, were imported into R Studio (v 3.6.1) and processed with custom scripts (https://github.com/ETHZ-INS/DLCAnalyzer). Values of points with low likelihood (> 0.95) were removed and interpolated using the R package “imputeTS” (v 2.7). The speed and acceleration of each point was determined by integrating the animals position over time. Points of interest relating to the arenas were tracked and median xy-coordinates were used to define the arenas *in silico*. The pixel-to-cm conversion ratio for each video was determined by comparing the volume of the arena *in-silico* in px^2^ to the measured size of the arena in cm^2^. Zones of interest were calculated from the arena definitions using polygon-scaling functions. We defined 6 zones in the OFT: the center (scaling factor = 0.5), the periphery (scaling factor = 0.8) and the 4 corners (scaling factor = 0.2, centered on the corners); and 3 zones in the EPM (open arms, closed arms and center). Integration of the body center over the entire video was used to calculate metrics such as total distance, average speed and time in zone for each mouse. Further, a speed cutoff (5 cm/s) was set to determine when and how long an animal was moving and its average speed whilst moving. Time floating in the FST was determined by analysing the rate of change of the polygon area formed by joining the head-centre, tailbase, bcl and bcr **(Figure 1)**. Whenever this rate of change (smoothed with a rolling mean over ±5 frames) was below a preset cutoff (15 px^2^/frame) the animal was considered floating. Head dips in the EPM were scored by observing if the ‘nose’ and ‘headcentre’ points were outside of the EPM arena.

### Time Resolved Skeleton Representation

A position and orientation invariant skeletal representation was created from the DLC tracked coordinates at each frame. The distances between pre-determined tracking point pairs, angles between pre-determined vector pairs and the areas of pre-determined polygons in each frame were calculated. The resulting skeletal representation contained a total of 10 distances, 6 angles and 4 areas. Additionally two boolean variables were included to check if the points (in this case nose and head center) were inside the arena or not.

This skeletal representation was used to create short sequences of the skeleton over pre-determined time intervals. In this case, an integration period of ±15 frames was chosen. The skeletal data from each of these intervals was then flattened into a longer skeleton-sequence-vector for each frame. The resulting skeleton-sequence-vectors were combined into a skeleton-sequence-matrix that describes a short sequence of the skeleton of ± 15 frames for each frame centered on the frame.

### Machine Learning Approach

In order to create a training data set, 20 videos of the OFT were manually labeled (using VIA video annotator [40]), indicating the onset and offset of selected behaviors. Labeled behaviors include ‘supported rear’, ‘unsupported rear’, and by default ‘none’. Videos were labeled by three independent raters. These sets of labeling data were used to train multiple neuronal networks for the classification of the selected behaviors (labelling data can be accessed here: https://github.com/ETHZ-INS/DLCAnalyzer/data. All videos can be found here: https://zenodo.org/record/3608658). For each labeled video the other 19 videos were used to train a neuronal network. The model was then cross-validated on the single video not included in the training set. This process was repeated with each raters labeling data, resulting in a total of 60 models and cross validations. The R package for tensorflow and keras were used for machine learning. Training and testing data were normalized within videos using a Z-score (x − mean(x) / sd(x)) method on non-boolean parameters. The training data was randomly shuffled before training. A sequential model with two hidden layers was trained (input shape: N = 682, L1: dense layer, N = 256, dropout rate = 0.4, activation = ‘relu’; L2: dense layer, N = 128, dropout rate = 0.3, activation = ‘relu’; and an output layer with: 4 nodes, activation = ‘softmax’). The network was trained for 10 epochs with a batch size of 32. The optimizer ‘rmsprop’, the loss function ‘categorical_crossentropy’ and metric ‘accuracy’ were used. Accuracy on the cross validation set was determined on a frame to frame basis. However, to remove single frame misclassifications the final classification was integrated over a period of ±5 frames.

### Statistical Analysis

Data was tested for normality and all comparisons between normally distributed datasets containing two independent groups were performed using unpaired t-tests (2 tailed) whereas all comparisons between more than two groups were performed using one-way ANOVAs in order to identify group effects. Significant main effects where then followed up with post hoc tests (tukey’s multiple comparison test). We also report the Coefficient of Variation (CV) in order to show the dispersion of the data around the mean.

## Results

Our goal was to compare the tracking performance of DLC to commercial solutions usingthree of the most popular rodent behavior tests in basic neuroscience research: the open field test, the elevated plus maze, and the forced swim test. Robust tracking was previously demonstrated using DLC [14] and other open source tracking software (e.g. ezTrack) [41], thus we established DLC tracking in arenas that are compatible with the commercial systems we routinely use in our lab. We labeled standardized points of interest when tracking the mouse in each test **(Figure 1a)**. The labels relating to the arenas are particularly important **(Figure 1b,c,d)**, as they enable the calculation of all of the standard parameters (e.g. time in center, distance travelled) from the frame-by-frame coordinates of each point of interest.

Next, we benchmarked DLC tracking performance against commercial behavioral tracking solutions. Where possible, we scored each test using the “tracking-only” software Ethovision XT14 (Noldus), and the “all-in-one” TSE Multi Conditioning system. We tested 20 mice in the TSE Multi Conditioning System’s OFT arena, the videos acquired from these tests were then analysed using EthoVision XT14 and DLC. In the OFT, simple tracking parameters such as distance travelled and time spent in zone (centre) were comparable between DLC and Ethovision. However, TSE’s Multi Conditioning system reported a significantly different mean distance travelled (One-way ANOVA, F(2,57)=331.9, P<0.0001, CV=DLC:12.24%, Ethovision:11.03%, TSE:16.83%), yet regarding time in zone reports a value similar to that of DLC and Ethovision (CV time in center= DLC:46.28%, Ethovision: 45.05%, TSE: 43.09%) **(Figure 2)**. Heatmaps can also be plotted from all systems showing that time in zone is for the most part comparable **(Figure S2)**. The vastly different distance reported by the TSE system is likely due to its reliance on an infrared beam grid, which predicts the centerpoint of the animal based on the number and location of the beams that are broken. Thus, slight movement of the animal can lead to relatively large movements of the centerpoint, which could inflate the total distance travelled. This issue does not appear to affect the time spent in zones, since the fluctuation of centerpoint is unlikely to be large enough to move across zones. The distance recorded by the TSE system also correlates poorly with the other systems, which might again be due to the confounding factors introduced by the beam grid (see discussion).

**Figure 2.**
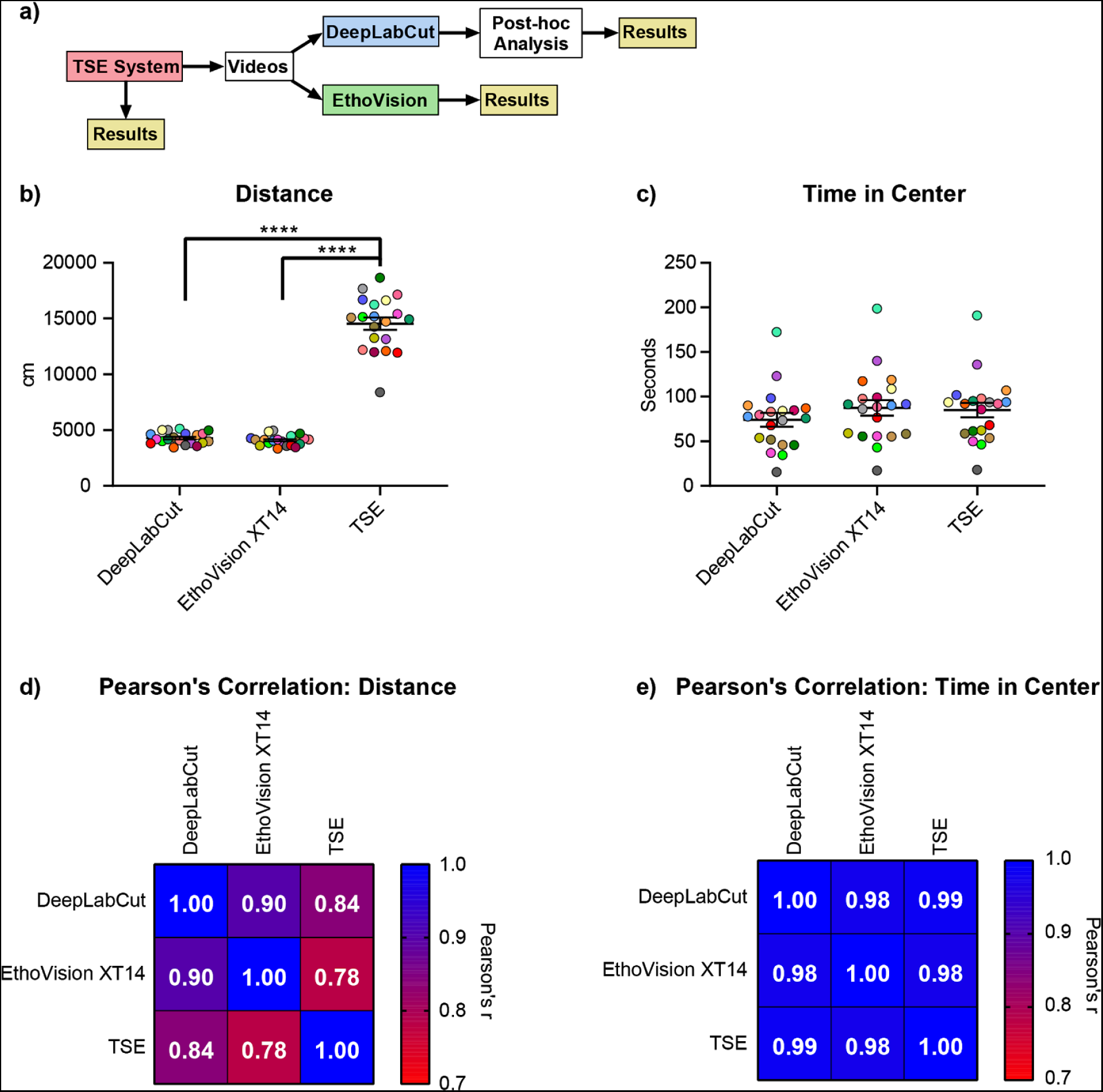
A comparison of basic tracking parameters in the open field test. **(a)** Schematic showing the workflow of the comparison between systems. **(b,c)** Distance and time in center as reported by DeepLabCut (with post hoc analysis), Ethovision XT14 and the TSE Multi Conditioning System (TSE). **(d,e)** Correlation analysis of the performance of the different systems. Data expressed as mean ± standard error of the mean. Colors represent individual animals and are consistent across analysis techniques for direct comparison (n=20) ****=p<0.0001.

The FST and EPM analyses could not be scored using the TSE Multi Conditioning System, since the EPM/FST apparatus is not compatible with its “all-in-one” setup. We therefore acquired videos of 29 mice performing the FST and 24 mice performing the EPM using EthoVision, which were later analysed using DLC. Using DLC and Ethovision XT14, we found no significant differences regarding distance travelled in the FST or EPM (CV distance in swim= 23.71% (DLC), 26.76%(Ethovision), CV distance in EPM= 32.75 (DLC), 32.74 (EthoVision)) or time in zones (CV Time in Open in EPM= 59.88 (DLC), 58.79 (EthoVision). CV Time in Closed in EPM= 37.66 (DLC), 36.26 (EthoVision). CV Time in Center EPM= 55 (DLC), 56.73 (EthoVision)) **(Figure 3)**.

**Figure 3.**
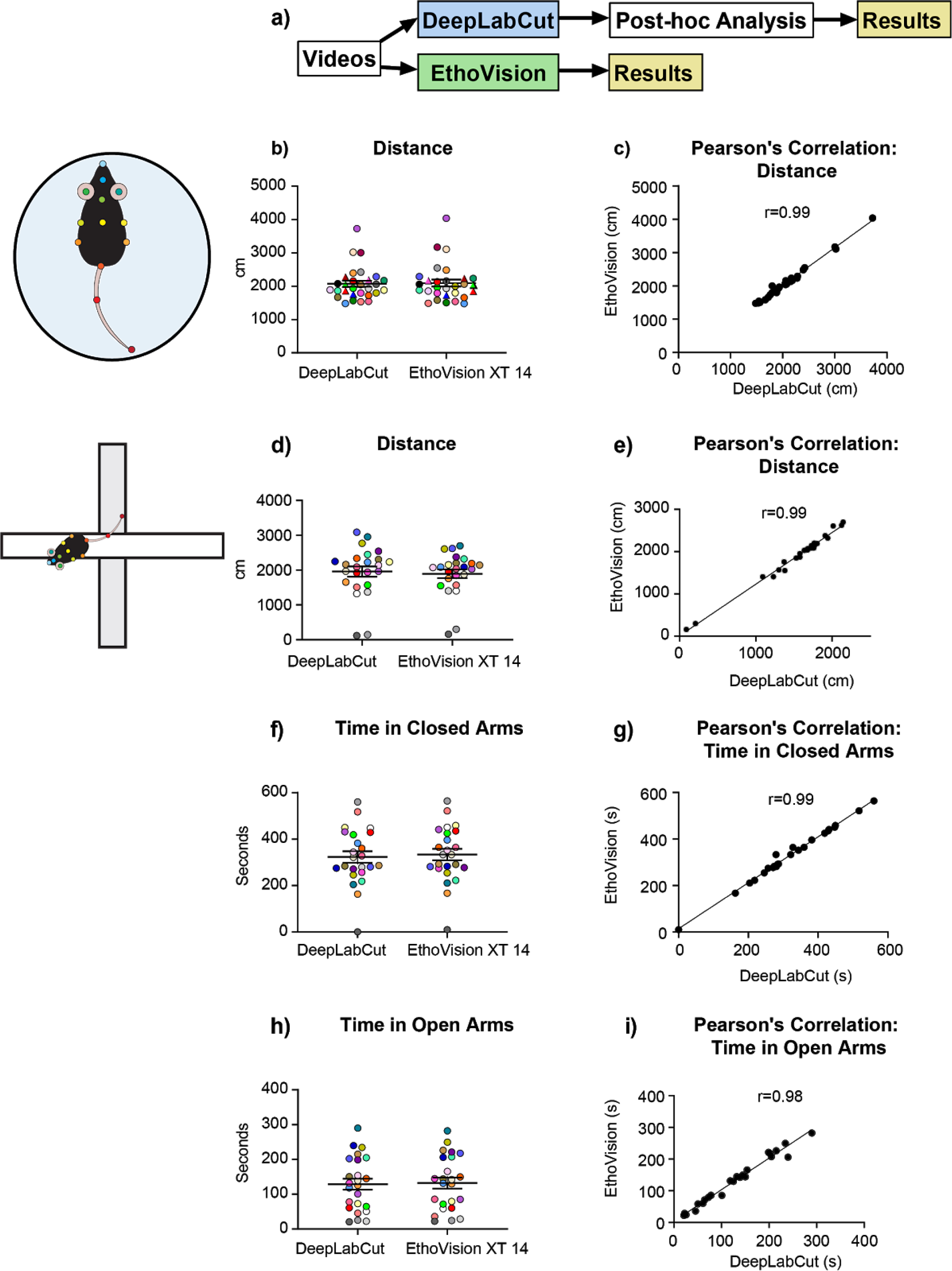
A comparison of basic tracking parameters in the forced swim test and elevated plus maze. **(a)** Schematic showing the workflow of the comparison between systems. **(b,d,f,h)** Basic tracking parameters in the forced swim test and elevated plus maze as reported by both DeepLabCut (with post hoc analysis) and Ethovision XT14. **(c,e,g,i)** Correlation between the scores of the two systems. Data expressed as mean ± standard error of the mean. Colors represent individual animals and are consistent across analysis techniques for comparison (FST n=29, EPM n=24) *=p<0.05.

Providing evidence that DLC can perform basic tracking functions similarly to commercial software/hardware packages, we next attempted to score ethological behaviors using the coordinates for each datapoint tracked by DLC. To establish the best possible ‘ground truth’, three human annotators manually scored floating behavior in a set of 10 forced swim test videos. Animals were considered to be floating if the rate of change of the polygon formed by joining the head-centre, tailbase, bcl and bcr **(Figure 1a and Figure 4b)** was less than 15 square pixels between frames. Using the same videos, we were able to accurately identify floating behavior **(Figure 4)**. Additionally, we compare this to the ‘activity’ module for Ethovision XT14, which can be used to score floating behavior **(Figure 4)**. We detected no significant differences in time floating, with EthoVision showing a better correlation with manual scoring than DLC. The differences between DLC and EthoVision are likely due to different levels of optimization and the different approaches to classifying movement, with EthoVision comparing changes at the pixel level frame to frame, and DLC only using the information gathered from the tracked coordinates. In the EPM, head dips were also recorded using DLC and Ethovision **(Figure 4)**. Here we saw significant group effects (One-way ANOVA, F(4,20) = 23.82, P < 0.0001, CV head dips= R1:50.71 %, R2:46.38 %, R3:45.54 %, DLC:47.03 %, Ethovision:28.33%), with differences between all groups and EthoVision (Tukey’s multiple comparisons test, q = 10.96(R1),11.50(R2),10.83(R3), 9.60 (DLC) df = 20, p < 0.0001), but no differences between human annotation and DLC. The difference in head dips is likely the result of different parametric definitions of the behavior which is addressed in more detail in the discussion.

**Figure 4.**
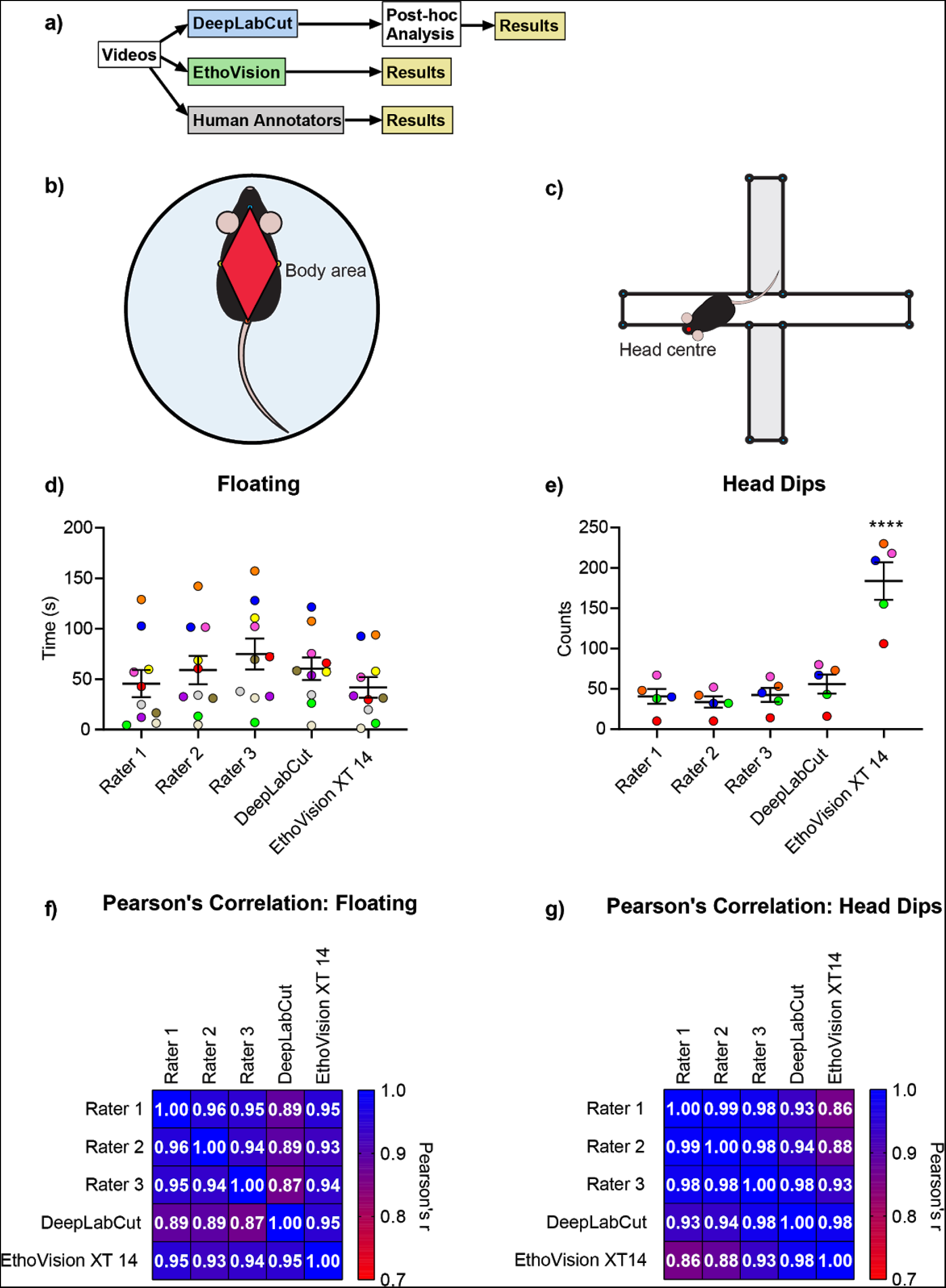
A comparison of floating and head dips, two basic ethological behaviors reported in the forced swim test/elevated plus maze. **(a)** Schematic of the workflow for the comparison between systems. **(b,c)** The polygon used in the definition of floating, and the points taken into account when defining head dips. **(d,e)** Floating in the forced swim test and head dips in the elevated plus maze as reported by 3 human annotators (rater 1-3), DeepLabCut (with post-hoc analysis), and Ethovision XT14’s behavioral recognition module. **(f,g)** Correlation analysis for comparison between approaches. Data expressed as mean ± standard error of the mean. Colors represent individual animals and are consistent across analysis techniques for comparison (FST n=10, EPM n=5) ****=p<0.0001.

So far, we have demonstrated that manually defined parameters can be used to automatically determine distinctive behaviors based on custom pre-defined criteria and simple post-hoc analysis of tracking data generated by DLC. However, we found that using this approach for more complex behaviors was labor intensive, arbitrary and inaccurate (data not shown). We therefore applied supervised machine learning to recognize complex behaviors in the open field test. We used the coordinates for each datapoint tracked by DLC to reconstruct a rotation and location invariant skeletal representation of the animal **(Figure S3)**. We then trained a small artificial neuronal network (2 layers, L1 = 256 neurons, L2 = 128 neurons, fully connected) to recognize short sequences of the skeletal representation during epochs of supported and unsupported rears. We focused on rearing in the open field since supported and unsupported rears are very similar movements (both include standing on hind legs), which are difficult to score automatically [29]. Again, we had 3 annotators scoring 20 videos (10 mins long) to set the ground truth for rearing frequency, and annotate the exact onset and offset of each behavior. We used the data of each annotator to train 20 behavior classifiers. To cross validate classification performance we trained each classifier on 19 videos and then tested on the remaining video. This allowed us to assess the classifier’s performance and to calculate correlation to the human annotation. Overall our behavior classifiers reached a frame-to-frame accuracy of 86 ± 3% **(Figure S4)**. No significant differences were observed between any of the human investigators (R1-3) or the machine learning classifiers trained using their data (MLR1-3) in the scoring of either supported rears (CV = 16.41 % (R1), 16.04 % (R2), 19.04 % (R3), 15.75 % (MLR1), 16.85 %(MLR2), 17.23 % (MLR3) or unsupported rears CV = 50.33 % (R1), 48.86 % (R2), 45.84 % (R3), 47.76 % (MLR1), 50.67 % (MLR2), 42.43 % (MLR3). Therefore, supported and unsupported rearing can be measured as accurately by supervised machine learning algorithms as by human manual scoring, the gold standard in the field **(Figure S5)**.

We then took the mean score from the human investigators and the mean score from the machine learning classifiers for each type of rearing and compared them to those reported by the TSE Multi Conditioning System, which includes a separate infrared tracking grid (z-grid, which counts beam-breaks as rears) and to Ethovision XT14’s behavior recognition module **(Figure 5)**. Significant group effects were observed in the scoring of unsupported rears (one-way ANOVA, F(3,76) = 9.547, p < 0.0001) with differences between the human raters and EthoVision (tukey’s multiple comparison test, q = 4.590, DF = 76, p = 0.0093), the machine learning based behavioral classifiers and Ethovision (tukey’s multiple comparison test, q = 6.841, DF = 76, p < 0.0001), and between EthoVision and TSE (tukey’s multiple comparison test, q = 6.213, DF 76, p = 0.0002). We observed significant group differences between the number of supported rears reported by Ethovision, TSE, and the human and machine learning classifiers (one-way ANOVA, F(3,76) = 104.5, p < 0.0001). Post-hoc tests reveal significant differences between the human raters and Ethovision (tukey’s multiple comparison test, q = 4.518, DF = 76, p = 0.0108), and between the human annotators and the TSE system (tukey’s multiple comparison test, q = 18.72, DF = 76, p < 0.0001). Similarly, the machine learning classifiers reported significantly different results to those reported by EthoVision (tukey’s multiple comparison test, q = 5.670), DF = 76, p=0.0008) and the TSE system (tukey’s multiple comparison test, q = 17.57, DF = 76, p<0.0001). The TSE system and Ethovision were also in disagreement (tukey’s multiple comparison test, q = 23.24, DF = 76, p < 0.0001). Again, no significant difference was detected between the performance of the humans or machine learning classifiers. In summary, we conclude that Ethovision reports an inaccurate number of unsupported rears, while both Ethovision and TSE perform very poorly on supported rears. It is important to note that we spent a considerable amount of time and effort calibrating the TSE system specifically to report unsupported rears accurately. However, it appears the TSE system cannot score both supported and unsupported rears accurately at the same time. In contrast, the supervised machine learning-based behavior classifiers performed as well as the human annotators, the gold standard in the field.

**Figure 5.**
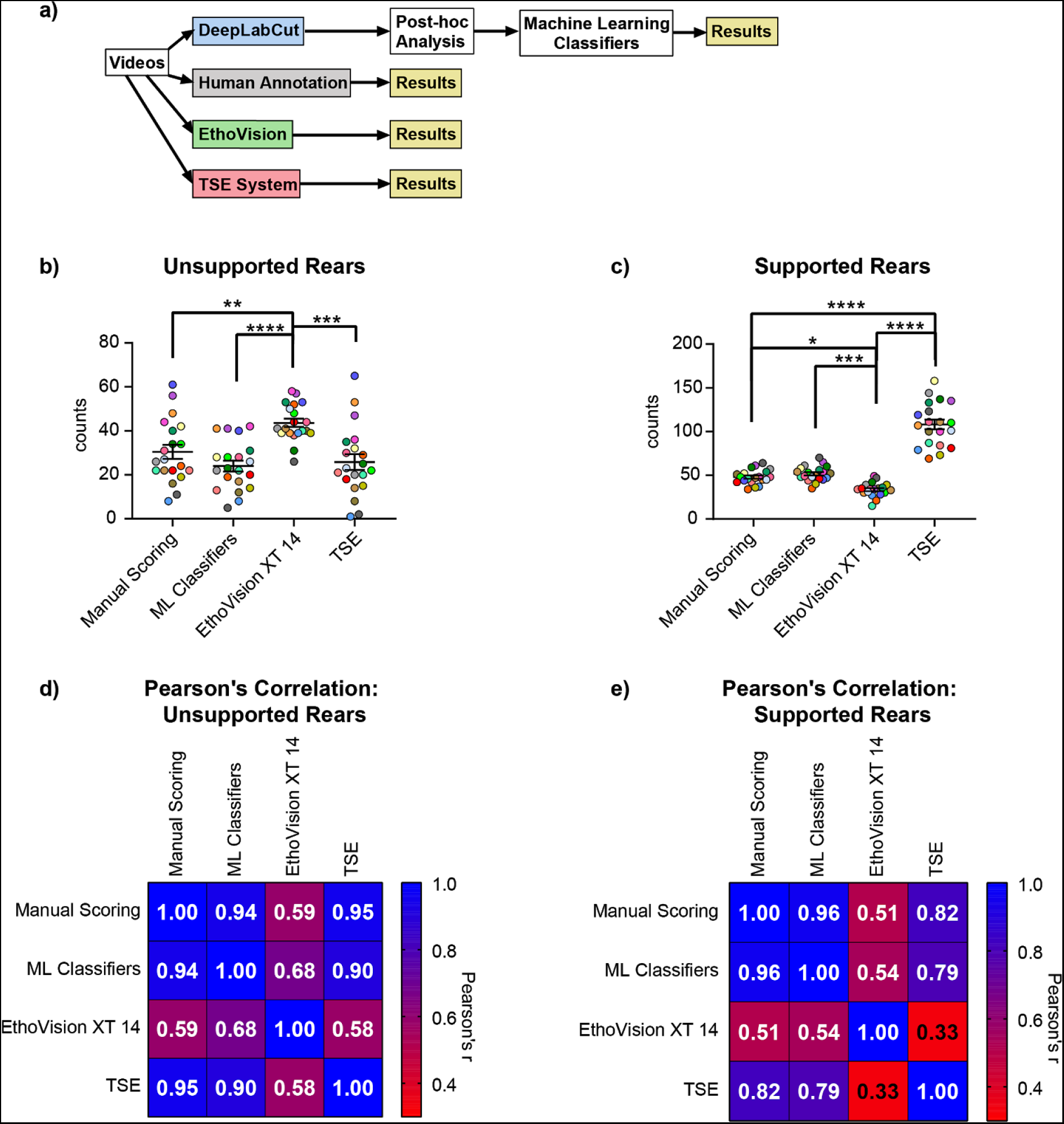
A comparison of complex behavioral scoring between human raters, machine learning classifiers and commercially available solutions. **(a)** Schematic of the workflow. **(b,c)** Unsupported and supported rears in the open field test as reported by 3 human raters (averaged and plotted as Manual Scoring) and 3 machine learning Classifiers (averaged and plotted as ML Classifiers), Ethovision XT14 and the TSE Multi Conditioning System (TSE). **(d,e)** Correlation analysis for comparison. Data expressed as mean ± standard error of the mean. Colors represent individual animals and are consistent across analysis techniques for comparison (n=20).*=p<0.05,**=p>0.01, ***=p<0.001,****=p>0.0001.

## Discussion

This report shows that DeepLabCut (DLC) video tracking combined with simple post-hoc analyses can analyse behavioral data as well as - if not better than - commercial solutions. With the addition of supervised machine learning approaches, the tracking data obtained from DLC can also be used to achieve close to human accuracy (the gold standard) when scoring complex, ethologically relevant behaviors, thus outperforming commercial systems at a fraction of the cost.

Scoring complex ethological behaviors with human-like accuracy is an important step forward in the analysis of behavior. Previous attempts to automatically score ethological behaviors have reduced intra-rater variability and increase throughput, but at the cost of accuracy. The machine learning classifiers used here are capable of reducing intra-rater variability by eliminating factors such as fatigue or human bias, whilst scoring accurately (similarly to a human investigator). This not only saves time but also money as similar approaches can be used to score any number of behaviors at no cost. Most importantly, within a given lab this approach allows consistent scoring with human accuracy, while avoiding inter-rater variability as experimenters change over time.

Regarding human scoring, our annotators were trained at the same time and reached a consensus about what constituted each behavior before beginning to score the videos. This likely reduced the inter-rater variability that can arise from differences in the definitions of the behaviors (even within a given lab). Additionally, the behaviors reported here were not scored in real time, which enabled frame-by-frame labelling. This offers advantages over live scoring, especially regarding fast or complex behaviors. Together, these factors likely explain why our inter-annotator scoring correlations are higher than some of those previously reported (approximately r=0.96 instead of r=0.90 for floating [8,42]). Although labelling post-hoc with this level of accuracy is time consuming (approximately 1 hour per 10-minute video), once DLC and the machine learning classifiers are trained, no further manual scoring is required, thus drastically reducing the overall time and effort required to accurately score behavior. The advantages of post-hoc labelling are passed on to the machine learning classifiers, which reproduce the higher quality labelling. In fact, machine learning classifiers often score animal behaviors more accurately than humans that often miss-label behaviors when scoring behavior in real time. In addition, humans often fail to score behaviors that occur in quick succession **(Figure S4)** for live v.s. post hoc behavioral scoring comparisons). We also tried to apply this machine learning approach to grooming behavior, however the low frequency of grooming events (approximately 20-30 events) in our training dataset led to insufficient training data and thus this behavior was excluded from our analysis. It is important to highlight that commercial systems allow altering the analysis parameters(to varying degrees). For the purposes of this report we tried to use the default/suggested settings where possible, and invested approximately equal amounts of time into the setup of all systems, thus giving a representable comparison whilst acknowledging that the performance of any system could still be improved (within limits).

The data presented here show that for simple parameters (such as distance travelled and time in zones), there appear to be large discrepancies between the values reported by infrared beam grids and video tracking software packages **(Figures 2,5)**. The exact distance travelled is not technically of importance in the majority of experiments, but it limits the ability to compare results across laboratories. Given the state of current technology there is no reason that this value should not be accurate, especially since the weaker correlation and wider spread of data gathered using the infrared beam grid could alter the interpretation of results and potentially mean that phenotypes go undetected. This is likely the result of the confounding factors of using a beam grid as opposed to video tracking (unspecific beam breaks/body center estimation techniques). Additionally, the number and spread of the beams determines precision and once the grid is built these features cannot be easily altered to fit the requirements of the user. Specialised infrared permeable arenas must also be used in combination with the beam grid, which further reduces the number of testing possibilities.

Video analysis packages such as Ethovision also have limitations, for instance Ethovision requires the test arena to be defined prior to analysis. Once the test arena has been defined this is no longer flexible, meaning that if the apparatus is moved slightly during cleaning it has to be returned to exactly where it was when the arena was defined. Although seemingly only a minor issue, this can drastically increase the amount of time required to score videos in which the camera/arena move slightly. Since DLC detects the arena, it is impervious to these slight movements and the calibration of the arena is always optimal regardless of the size of objects in the video, making it less prone to errors when the setup is used for multiple tests. DLC could also prove useful when working under more ethological conditions in arenas with bedding material/variable backgrounds, where commercial solutions will likely struggle even more, while the power of the deep learning approaches will get to flex their muscles.

Another key advantage of using DeepLabCut is that it offers increased tracking flexibility, which enables the user to define and record the exact parameters they are interested in, without the unnecessary constraints or paywalls many commercial systems have in place. For example, analyzing “head dips” or “rearing” requires the purchase of an additional module from EthoVision. Additionally, by defining each feature, it is also possible to know exactly what is being measured. Commercial packages often give explanations as to how they define their parameters, but often these definitions are not consistent between different commercial solutions and cannot be altered. This difference in definitions is the reason that EthoVision scores head dips so poorly in comparison to human investigators **(Figure 4)**. It would appear that EthoVision defines a head dip as the entry of the nose into the head dip area, which is located just outside of the open arm zones. The movement of the nose into the ‘head dip area’ is a prerequisite for a head dip, but not every movement of the nose into this zone will result in a head dip. Most human observers only score a head dip when the animal actually dips its head over the side of the maze as though the animal is looking at the floor. This discrepancy means that human observers will be reporting a completely different behavior. Moreover, the TSE system’s definition of rearing is also less flexible than one that can be generated using the tracking data from DLC. As the TSE system relies on an infrared beam grid it cannot distinguish between different behaviors that may break these beams and is therefore inaccurate. As seen in **Figure 5**, it can be adjusted to give an accurate account of unsupported rearing, but this comes at the cost of being able to identify other behaviors, in this case supported rearing. Other groups may also be interested in different behaviors such as stretching or grooming. EthoVision can detect these behaviors with some degree of accuracy [43], after the behavioral recognition module has been purchased. The TSE Multi Conditioning System cannot detect either of these behaviors as they are difficult to define when using an infrared beam grid for tracking. DLC could be used to identify any of these behaviors with either simple multipoint tracking (to detect stretching), or by applying a machine learning approach like the one above used to detect rears (for more complex behaviors such as grooming).

As behavioral analysis moves more toward video tracking as opposed to reliance on beam grids, recent developments in unsupervised behavioral identification approaches have widened the horizons of what was previously thought possible. Approaches that focus on the unsupervised identification and separation of behavioral patterns are beginning to reveal the true complexity and richness of animal behavior [9,10,13], However the interpretation of the findings from unsupervised machine learning techniques are more difficult. Although impressive, the implementation and use of many of these unsupervised behavior recognition approaches is out of reach of many basic science labs that lack the necessary programming and machine learning know-how. Therefore, widespread use/dissemination of new cutting-edge techniques will likely depend on their commercialization as part of user-friendly software/hardware solutions. In contrast, modern deep learning/machine vision based tracking and behavioral identification approaches such as those demonstrated here using DeepLabCut, are already taking over the field of behavioral neuroscience. In this first systematic, head-to-head comparison, we show that they are ready to be deployed in the field, offering high accuracy and precision while being flexible and affordable.

## Acknowledgments

We would like to thank Rebecca Waag, Sian Duss and Jin Qiuhan for their help scoring animal behaviors and Jens Weissman, Roger Staub and Sanja Vasic for looking after the animals used in these experiments. We also thank Mackenzie Mathis for critical reading of the manuscript.

## Author Contributions

Conceptualization, OS, LVZ, JB; Methodology, OS, LvZ, CS, FA; Investigation, OS, CS, FA; Writing – Original Draft, OS, LvZ, JB; Writing – Review & Editing, OS, LvZ, BG, JB; Funding Acquisition, JB, BFG; Resources, JB; Supervision, OS, LvZ, BFG, JB.

