## Supplementary material for "Deep learning based behavioral analysis enables high precision rodent tracking and is capable of outperforming commercial solutions": Figure S1

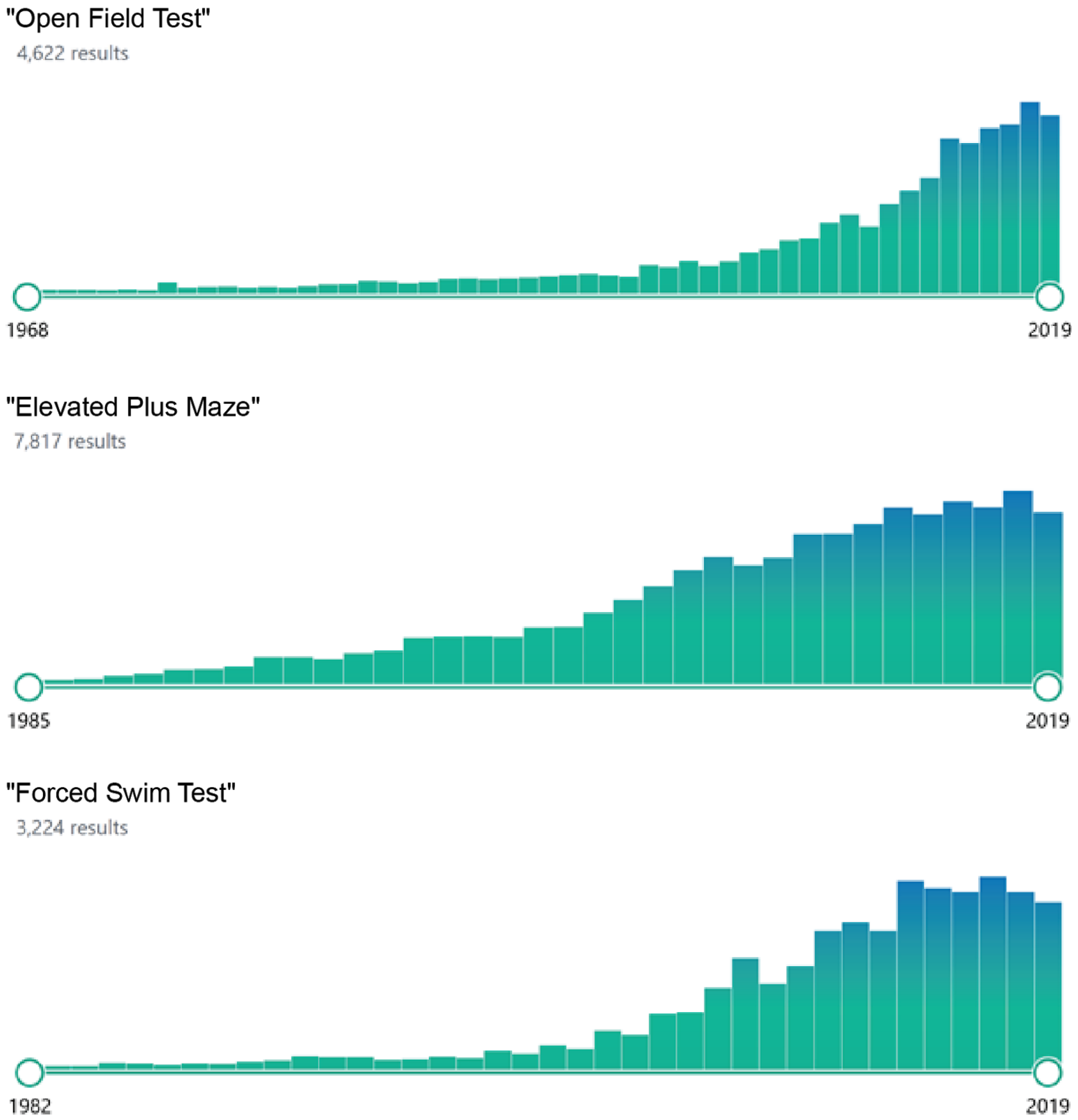


**Figure S1. The number of citations relating to the Open Field Test, Elevated Plus Maze and Forced Swim Test.** Over the last few decades the popularity of these behavioral assays has increased dramatically, making them three of the most popular behavioral assays used in behavioral neuroscience.
