## Supplementary material for "Deep learning based behavioral analysis enables high precision rodent tracking and is capable of outperforming commercial solutions": Figure S2

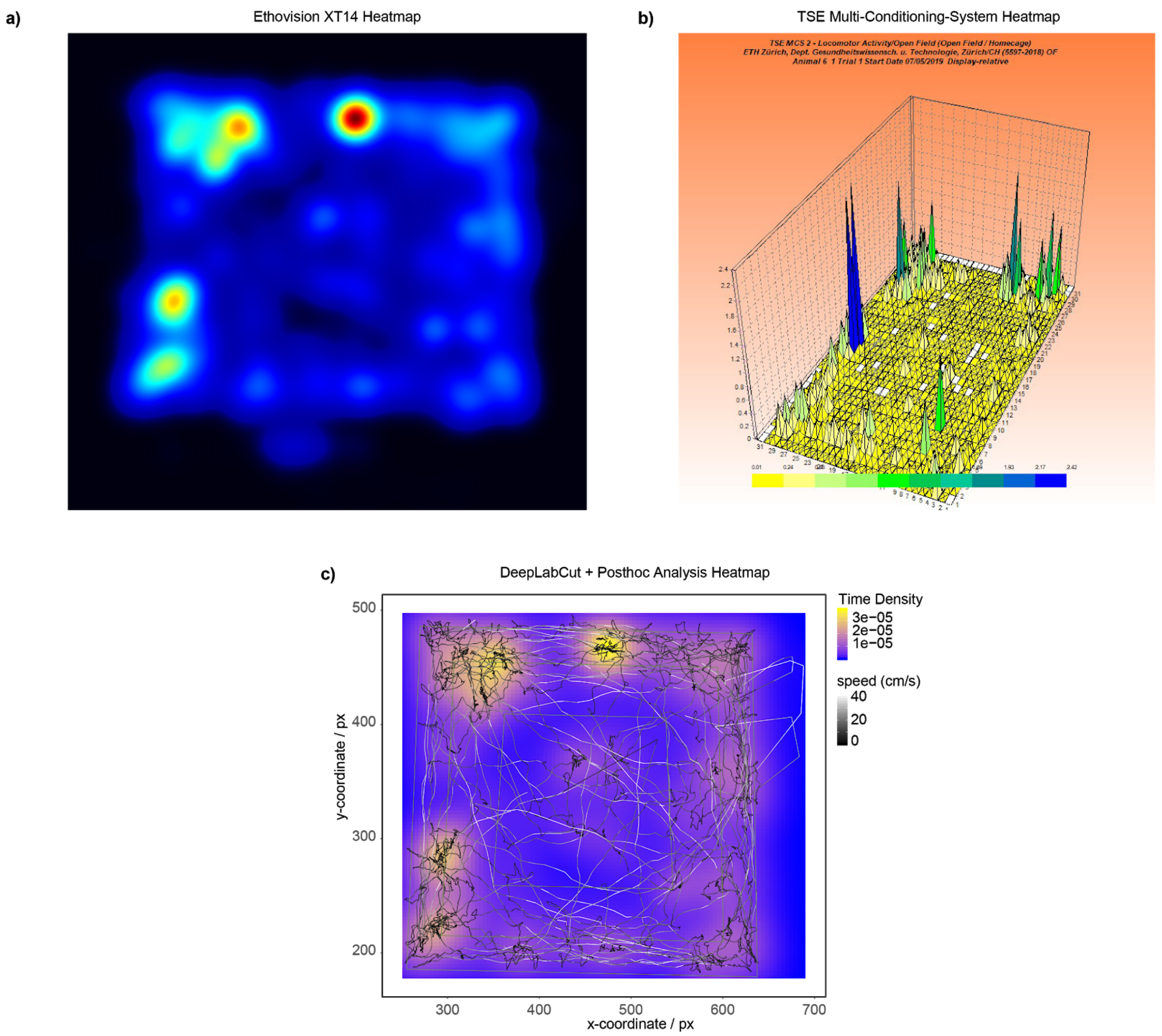


**Figure S2. Heatmaps.** example heatmaps generated using (a) EthoVision XT14, (b) TSE's Multi Conditioning System and (c) DeepLabCut (with post hoc analysis and ggplot2).
