## Supplementary material for "Deep learning based behavioral analysis enables high precision rodent tracking and is capable of outperforming commercial solutions": Figure S3

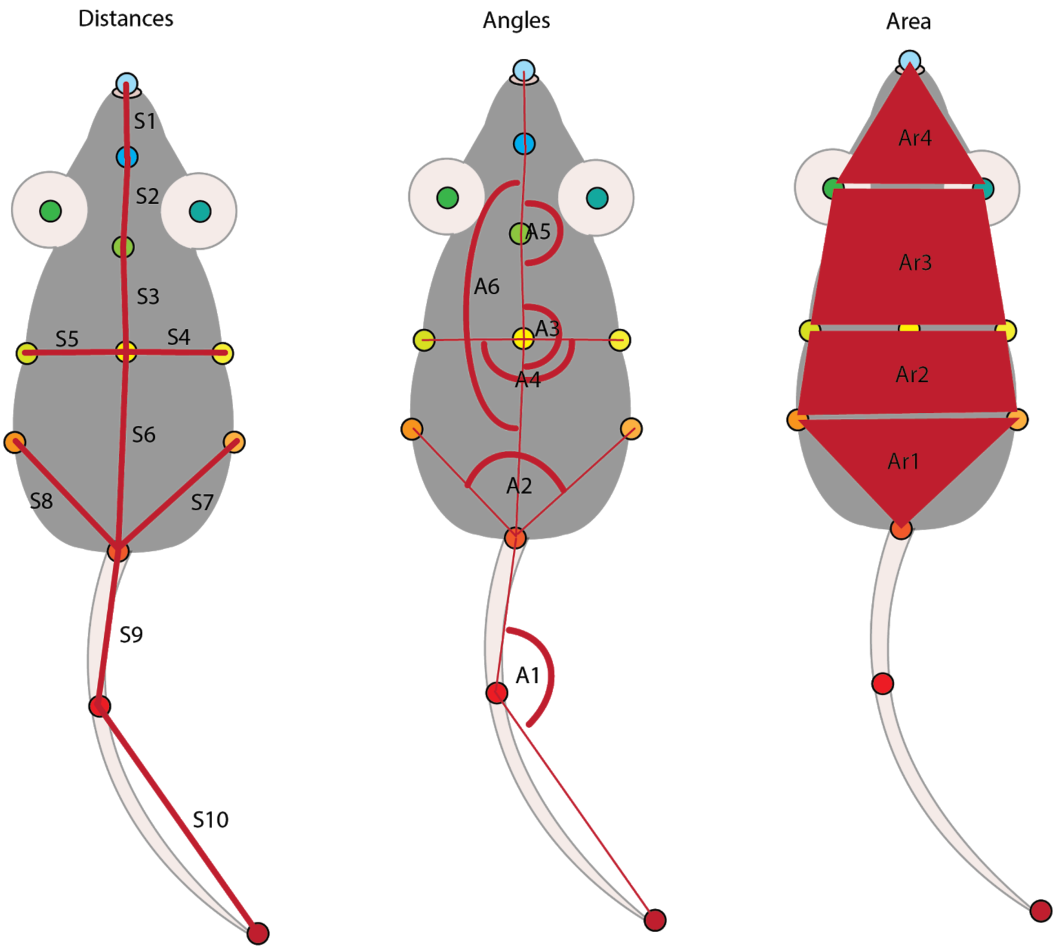


**Figure S3. Skeletal data for machine learning.** The skeletal information taken into account during the machine learning process, gathered from the tracking data obtained using DeepLabCut.
