## Supplementary material for "Deep learning based behavioral analysis enables high precision rodent tracking and is capable of outperforming commercial solutions": Figure S4

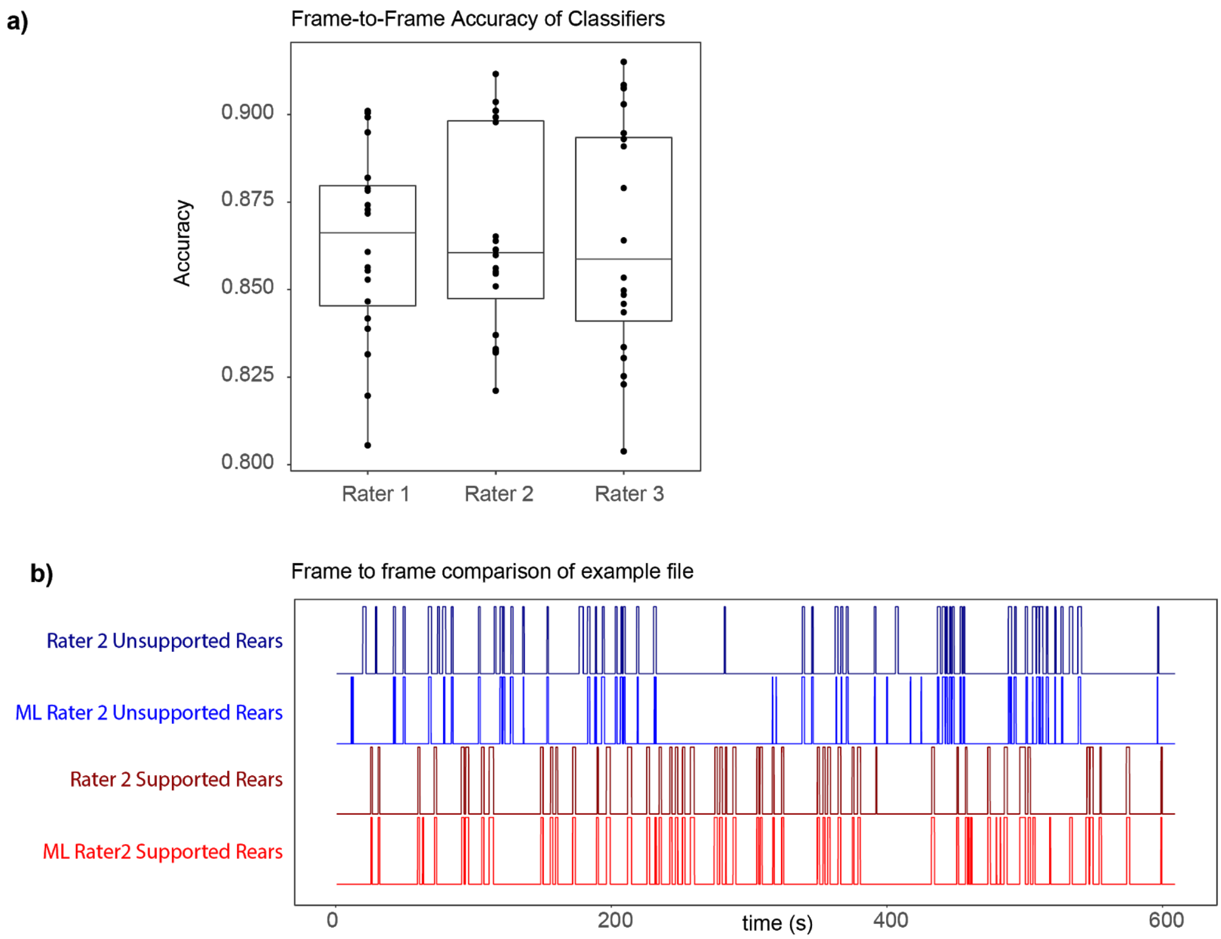


**Figure S4. Frame by frame comparison of scoring by a single human rater and their machine learning counterpart**. (a) the frame by frame accuracies of each rater and the behavioral classifier trained using their labelling data. (b) an example of the frame by frame labelling, each bar indicates the onset and offset of the indicated behavior. Although some inconsistencies can be observed, the total number of behaviors reported is similar with a frame-to-frame accuracy of 86 ± 3%.
