## Supplementary material for "Deep learning based behavioral analysis enables high precision rodent tracking and is capable of outperforming commercial solutions": Figure S5

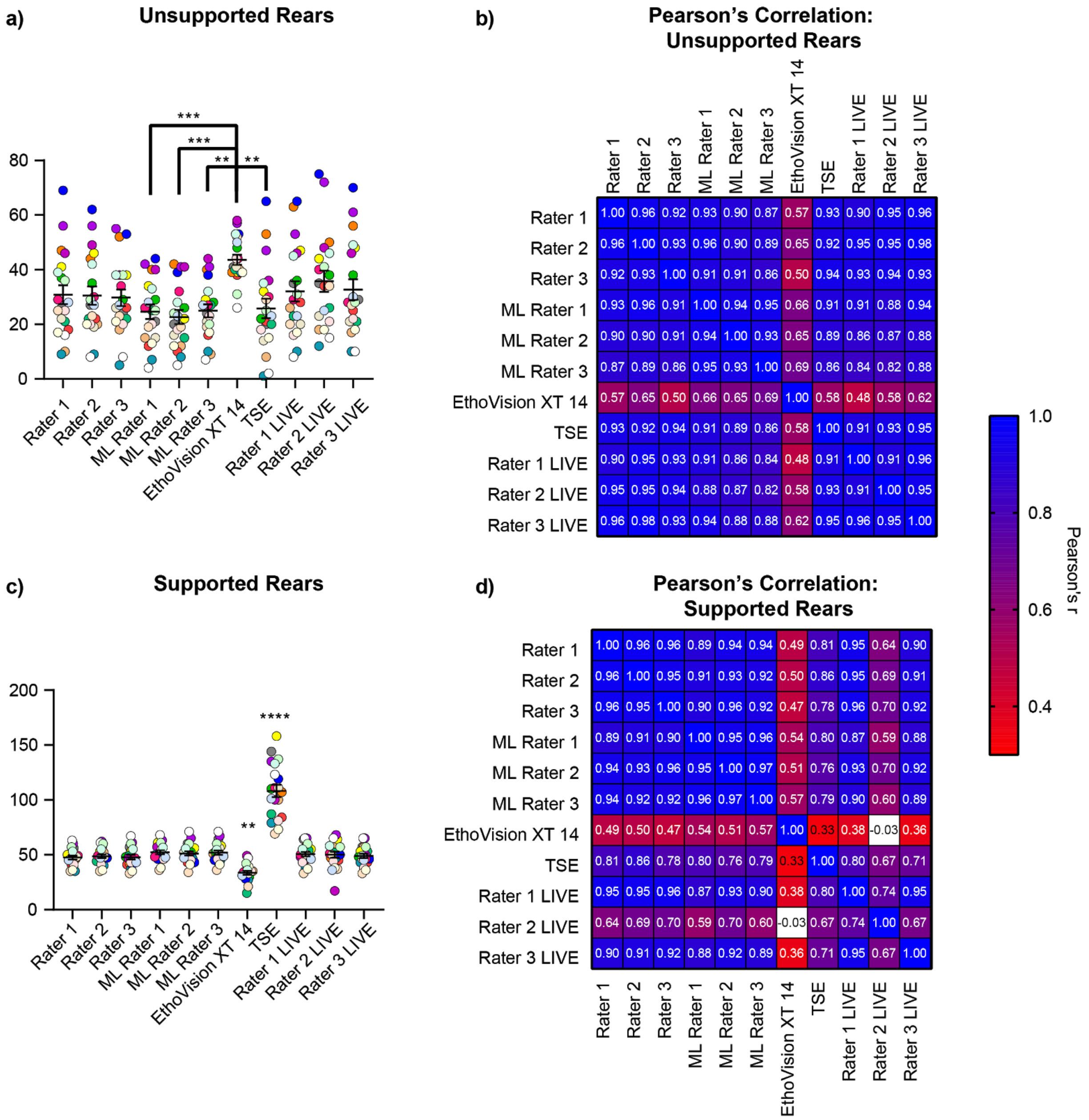


**Figure S5. An expanded version of Figure 5.** Absolute values of supported and unsupported rears in the OFT, and correlation plots to compare different scoring techniques. (a,c) The number of unsupported (a) and supported (c) rears reported by the three human annotators (rater 1-3), the three machine learning classifiers generated using the training data from the human annotators (ML rater 1-3), EthoVision XT14, the TSE Multi Conditioning system (TSE), and additionally the live scoring from the human annotators. (b,d) Correlation analysis comparing different scoring techniques. All raters and ML raters were significantly different to both EthoVision XT14 and TSE, only the highest p-values are reported for clarity. Colors represent individual animals and are consistent across analysis techniques for comparison (n=20),**=p>0.01, ***=p<0.001,****=p>0.0001.
